# Empty sella syndrome as a window into the neuroprotective effects of prolactin

**DOI:** 10.1101/2020.11.30.403576

**Authors:** David A. Paul, Emma Strawderman, Alejandra Rodriguez, Ricky Hoang, Colleen L. Schneider, Sam Haber, Benjamin L. Chernoff, Ismat Shafiq, Zoë R. Williams, G. Edward Vates, Bradford Z. Mahon

## Abstract

**BACKGROUND:** To correlate structural integrity of visual pathway white matter tracts with prolactin levels in a patient who demonstrates downward herniation of the optic chiasm secondary to medical treatment of a prolactinoma.

**METHODS:** A 36-year-old woman with a prolactinoma presented with progressive bilateral visual field defects nine years after initial diagnosis and medical treatment. She was diagnosed with empty-sella syndrome and instructed to stop cabergoline. Hormone testing was conducted in tandem with routine clinical evaluations over one year and the patient was followed with diffusion magnetic resonance imaging (dMRI), optical coherence tomography (OCT), and automated perimetry at three time points. Five healthy controls underwent a complementary battery of clinical and neuroimaging tests at a single time point.

**RESULTS:** Shortly after discontinuing cabergoline, diffusion metrics in the optic tracts were within the range of values observed in healthy controls. However, following a brief period where the patient resumed cabergoline (of her own volition), there was a decrease in serum prolactin with a corresponding decrease in visual ability and increase in radial diffusivity (p<0.001). Those measures again returned to their baseline ranges after discontinuing cabergoline a second time.

**CONCLUSIONS:** These results demonstrate the sensitivity of dMRI to detect rapid and functionally significant microstructural changes in white matter tracts secondary to alterations in serum prolactin levels. The inverse relations between prolactin and measures of diffusion and visual function provide support for a neuroprotective role of prolactin in the injured nervous system.

## INTRODUCTION

Crush and stretch injuries to the optic nerve, tract and chiasm in animals reveals that demyelination is a primary process in the progression of delayed axonal degeneration (1–4). Human studies looking at compression of the optic chiasm have confirmed these findings and also suggested the possibility of rapid vision recovery (5, 6). Unlike primary axotomy, in which axonal connections are immediately severed, delayed axonal degeneration is a potentially reversible process. Several mechanisms likely contribute to recovery, including remyelination (7), microtubule reorganization, shifts in ion channel permeability and restoration of glial-neuronal connections at the paranodes (8). Left unhindered, delayed axonal degeneration and its associated sequela lead to irreversible cell death (1–3, 9–11). Prolactin, a hormone synthesized in and released by the anterior pituitary gland with well-established roles in lactation, demonstrates promise as a mediator of delayed axonal degeneration and is known to control important mechanisms in the central nervous system (12, 13). In the retina, prolactin serves as a neurotrophic factor required for maintaining homeostasis during both injury (14) and physiologic aging (15), with receptors prominent in the ganglion cell layer as well as the outer and inner nuclear layers (16). In white matter, high prolactin levels signal oligodendrocyte proliferation, modulate neurotransmission (17) and promote white matter repair after demyelination in mice (18).

A specific disease model – compression of retinofugal fibers by large prolactin-secreting pituitary tumors (i.e. prolactinomas) provides a window with which to observe the potential mechanisms through which prolactin exerts a neuroprotective role in the injured human brain (19). Patients with prolactinomas often experience a stereotyped loss of vision in the temporal hemifields secondary to mass effect, in addition to multiple endocrinopathies (20). Treatment using dopamine agonists (e.g. cabergoline), increases the available substrate capable of binding D2 receptors on lactotroph cells in the anterior pituitary gland and inhibits prolactin release (21). A secondary effect of cabergoline is reduction in tumor size (21). As such, it is used as a first line treatment for prolactinomas. Surgical decompression is reserved for cases of dopamine agonist-resistant prolactinomas or acute changes in tumor size that necessitate immediate intervention to prevent blindness. Dopamine agonist therapy for large prolactinomas can rarely cause symptomatic empty-sella syndrome. A syndrome characterized by both downward herniation of the optic chiasm into an empty sella turcica (the bony cave that houses the pituitary gland) and delayed secondary vision loss. Only 21 cases are known to be reported in the literature, a large majority of them treated with either reduction or cessation of dopamine agonist therapy (22–27), or with surgical management to untether or elevate the optic apparatus (28–34). With cessation of dopamine agonist therapy, all reported patients demonstrated improvement of visual outcomes without any changes in either the level of herniation or appearance of the optic chiasm. How vision returns despite persistent downward herniation of the chiasm in this small cohort of patients has not previously been studied.

Here we investigate mechanistic hypotheses of the effect of prolactin on glial-neuronal function in an observational study of a rare patient with symptomatic empty sella syndrome. Both delayed axonal injury and retinal health were studied longitudinally at three time points over a year, using a combination of diffusion magnetic resonance imaging (dMRI) and optical coherence tomography (OCT), respectively – and compared with a cohort of healthy control participants. Indexed measures of diffusion have been used as an in vivo proxy for myelin integrity (3, 35–38). Specifically, an increase in radial diffusivity (RD) without changes in axial diffusivity (AD) indicates a breakdown in myelin, while decreased AD indicates axonal degeneration (37). These properties have been leveraged in human subjects to characterize microstructural changes occurring across the length of the optic tract for numerous pathologies that include pituitary macroadenomas (5), optic neuritis (39), and glaucoma (40). Additionally, retrograde degeneration of retinal ganglion cells can be characterized as a function of retinal nerve fiber layer (RNFL) and ganglion cell complex (GCC) thickness as measured by OCT (41–43).

Notably, the patient under study temporarily resumed cabergoline on her own volition prior to her second research visit. In retrospect, that event provided a natural contrast for observing the effects of varying levels of serum prolactin on measures of visual function, white matter integrity, and retinal health that otherwise would not be possible to observe. White matter integrity and retinal health were correlated with both serum levels of prolactin and ophthalmologic assessment of visual function (Humphrey perimetry) – allowing us to infer a causal relation between prolactin and maintenance of the coupled structure-function relationship between white matter integrity and visual ability. Secondly, this study demonstrates that diffusion MRI metrics are sensitive to detect early changes in white matter associated with varying levels of serum prolactin.

## METHODS

### Participant Recruitment

This research was conducted as part of an ongoing pituitary tumor research study approved by the Research Subjects Review Board at the University of Rochester (RSRB00071763). Patient AJ, a 36-year old woman with a large prolactinoma was recruited as part of this study. See supplemental text, S1 for a detailed clinical history. Five healthy control participants (n = 10 total hemifields, mean age 35.8 ± 11.88 SD) were also recruited and reviewed by an ophthalmologist (ZW) to confirm eligibility. Exclusion criteria included glaucoma, diabetic retinopathy, history of central retinal artery occlusion, optic disc drusen, multiple sclerosis, stroke, and previous head trauma. Supplemental Table 1 displays basic demographic information for all study participants. All participants gave written consent for participation in the study.

### Measurement of Serum Prolactin Levels

Serum prolactin was obtained via laboratory blood draw by a trained phlebotomist and analyzed by the University of Rochester Clinical Laboratories as part of the patient’s routine clinical care. The prolactin was measured via Electrochemiluminescence Immunoassay with a reference range of 4.8-23.3ng/ml. Approximately 10 laboratory draws were performed over the course of a year (see supplemental Figure 1) and data from each research visit was paired with the closest obtained laboratory value for analysis. Control participants each had a one-time blood draw obtained at a single University of Rochester Medical Center outpatient clinic on the same date as their OCT or dMRI testing, and analyzed by the same clinical laboratory as patient, AJ.

### Ophthalmologic Evaluation

#### Formal Ophthalmologic Testing

Automated 24–2 Humphrey perimetry (Zeiss HFA II-i series) and three-dimensional macular cube OCT (Zeiss Cirrus HD-OCT model 5000, 512×128 scan protocol with 6×6×2mm volumes) were performed for each eye – right eye (OD) and left eye (OS). Peripapillary retinal nerve fiber layer thickness (pRNFL) was additionally obtained using an optic disc cube 200 × 200 protocol. Testing was performed three times over a year for patient AJ, and at a single time point for all control participants. Crawford and Howell’s modified t-test was used to compare retinal thickness measurements and mean deviation for each eye at each time point for patient AJ to the control population (44).

#### Ganglion Cell Complex Thickness and Hemiretina Data

Macular central subfield (CST), ganglion cell layer (GCL), inner plexiform layer (IPL) and pRNFL thickness measurements were processed independently by Carl Zeiss Meditec, Inc. Ganglion cell complex (GCC) thickness was calculated using the sum of GCL and IPL layers. Measures of retinal thickness were subsequently mapped onto visual space using an in-house pipeline implemented in MATLAB (45) and analyzed with respect to visual hemi-field (i.e. the averaged thickness of homonymous hemiretinas) for comparison with optic tract integrity. To account for contributions from nasal versus temporal halves, retinal measures were weighted 53% contribution from nasal hemi-retina and 47% for temporal hemi-retina (46). The relations between weighted retinal thickness by hemi-field and corresponding optic tract diffusion metrics across all participants were evaluated using linear regression. Processed OCT data from Zeiss was unavailable for one healthy control due to acquisition artifact and thus excluded from the GCC analyses (controls n = 8 total hemifields).

### Magnetic Resonance Imaging Acquisition and Processing

#### MRI Acquisition and Analysis

Scanning was performed at the University of Rochester Center for Advanced Brain Imaging and Neurophysiology on a 3T Siemens MAGNETOM Prisma scanner with a 64-channel head coil. T1 weighted images were acquired at the start of each session with a MPRAGE pulse sequence (TR=2530 ms, TE=3.44 ms, flip angle=71°, FOV=256×256 sq mm, matrix =256×256, resolution=1 cu mm, 192 sagittal slices). Diffusion MRI data were acquired using a single-shot echo-planar sequence (65 diffusion directions, echo spacing=0.66ms, EPI factor=172, b=0, 1000, 3000s/sq mm, 96 slices, resolution=1.5 cu mm, 68 non-diffusion weighted volumes). Three non-diffusion weighted volumes were collected at the same resolution with reversed phase-encode blips to estimate the susceptibility-induced off-resonance field as implemented in FMRIB software library, or FSL (47, 48). FSL utilities were used to reduce motion artifacts and eddy current distortions and perform brain extraction (49). Probabilistic tractography of the optic tracts was performed on the preprocessed b=1000 files, using two fibers per voxel and Bayesian estimation (50–52), following techniques previously described (5, 53). All data in the main text are reported at a threshold of 2% for radial (RD) and axial (AD) diffusivity. See Supplemental Text, S2 for additional discussion on tractography, threshold determination, and supplemental analyses related to fractional anisotropy and mean diffusivity.

### Structural Analysis of the Optic Tracts

Optic tract cross-sectional area (CSA) was approximated using T1-MPRAGE scans of both patient AJ and controls, assuming an elliptical shape. Measurements were made using Horos, an open source DICOM viewer, freely available for download at: www.horosproject.org. Height and width were evaluated just posterior to the optic chiasm. One trained researcher (RH) obtained measurements for all participants. A Welch’s t-test was done to compare right and left optic tract CSA of this study’s controls with the respective CSA measured by Andrews et al., 1997, with no statistically significant difference between the data sets (54). The Crawford and Howell’s modified t-test was subsequently used to compare the patient’s right and left optic tract CSA at each time point with the healthy controls (44). Linear regression analyses were used to relate CSA with prolactin and diffusion MRI metrics.

## RESULTS

### Clinical Presentation: Patient AJ, a 36-year-old Woman with Empty Sella Syndrome

Patient AJ presented to the UR Medicine Pituitary Program at the University of Rochester with complaints of progressive bilateral visual field defects, photosensitivity and bilateral ocular pain nine years after initial diagnosis and subsequent medical treatment of a large prolactin-secreting pituitary tumor (Figures 1A-B). Clinical MRI demonstrated both a significant reduction in tumor size compared with her initial diagnosis (9 years earlier) and downward herniation of the chiasm into an empty sella turcica (Figure 1C). Ophthalmologic examination revealed bitemporal visual field defects on Humphrey perimetry (AJ0, Figure 1D) with intact visual acuity (20/20 in both OS and OD) and normal pRNFL (OD, 75 μm; OS 88 μm) (AJ0, Figure 1E). Prolactin levels were measured at 17.9 ng/mL, consistent with continued use of cabergoline. In keeping with previously published treatment recommendations (22–27), AJ’s cabergoline dose was reduced to 0.25 mg every other week and eventually discontinued. She was enrolled in the current study and serum prolactin and cabergoline dosage continued to be monitored as standard of care. At AJ’s first research visit (AJ1), her prolactin level was 86.6 ng/ml (normal reference range 4.8 – 23.3 ng/ml for non-pregnant females at our institution), consistent with treatment recommendations. Her prolactin level then dropped to 23.5 ng/ml after she resumed taking cabergoline on her own volition at the second time point (AJ2), and subsequently rose to 179 ng/ml at the third time point (AJ3) after discontinuing cabergoline a second time. These clinical data confirm already established relations between cabergoline use and serum prolactin. See Supplemental Figure 1 for a historical timeline of serum prolactin levels as a function of cabergoline dose and supplemental text S1 for a detailed clinical history.

**Fig. 1.**
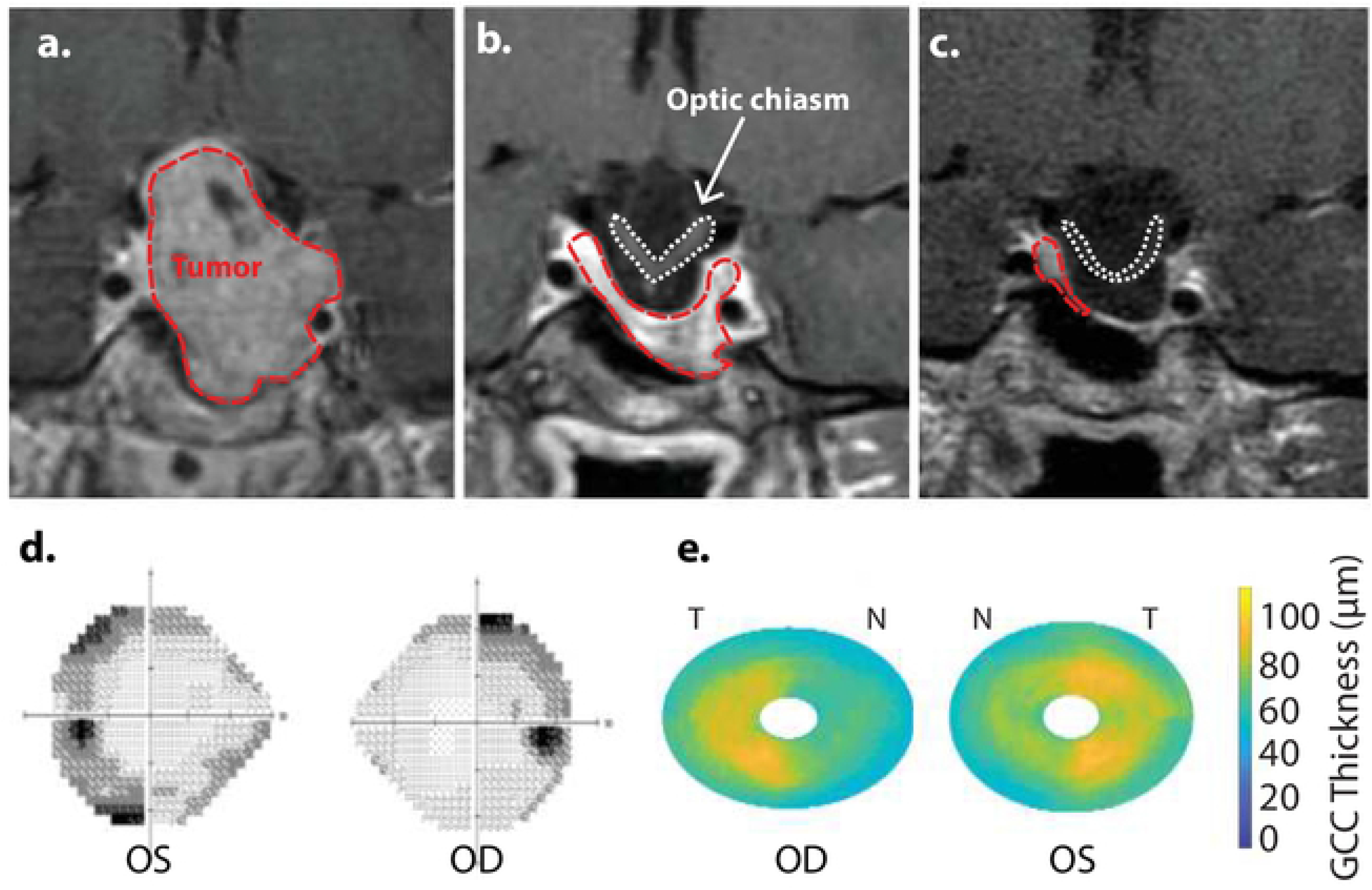
Clinical T1-weighted MRI scan with contrast demonstrating patient AJ’s pituitary macroadenoma in 2009 prior to (a), and after initiation of treatment with cabergoline (b), with progressive herniation of the optic chiasm into an empty sella (c) that was present at the time of study enrollment in 2018. Initial imaging of the mass (a) demonstrated significant contrast enhancement with expansion of the sella-turcica, asymptomatic extension of the tumor into the left cavernous sinus, and suprasellar extension with some deviation, but no outright compression of the optic chiasm. The normal pituitary gland and stalk were pushed to the right and superiorly. (d) Humphrey 24-2 visual field automated perimetry scan in 2018 at the time of empty sella diagnosis and prior to discontinuation of cabergoline and study enrollment, with OD mean deviation, −7.63 dB and OS mean deviation, −10.05 dB. (e) GCC thickness map obtained at study onset with nasal (N) and temporal (T) halves labelled according to the associated visual hemifield (e.g. OS nasal hemiretina represents the OS temporal visual hemi-field, T). Reduced thickness is primarily in the nasal hemiretina bilaterally.

### Ophthalmologic Evaluation of Visual Function and Retinal Thickness

#### Visual Function

At all three time points our patient demonstrated significantly reduced visual function compared with healthy control participants, as measured by mean deviation (all p < 0.05, using Crawford and Howell’s modified t-test; Table 1). This difference is most pronounced at AJ2 relative to the other time points and overlaps with the period in which the patient resumed cabergoline. Figures 2A-C display Humphrey 24-2 visual automated perimetry data for patient AJ at each time point, demonstrating a transient worsening of vision while on cabergoline.

**Table 1.** Mean deviation in AJ compared with healthy controls. Demonstrated here is the average perimetric mean deviation (in decibels, dB) for both healthy control participants (n = 10 total eyes) and patient AJ across all time points, with AJ0 representing visual field data acquired on initial presentation just prior to study enrollment. Error measurements for the cohort of healthy controls are reported as standard deviation; * p < 0.05, † p < 0.01. Refer to Supplemental Tables 4-5 for individual mean deviation values of all participants in the study reported both by eye and hemiretina.

|  | **Table 1. Mean deviation in AJ compared with healthy controls.** |  |  |  |  |
| --- | --- | --- | --- | --- | --- |
|  | Healthy Controls | AJ0 | AJ1 | AJ2 | AJ 3 |
| OD Mean Deviation (dB) | -1.99 ± 0.69 | -7.63<sup>†</sup> | -6.57<sup>†</sup> | -9.51<sup>†</sup> | -9.65<sup>†</sup> |
| OS Mean Deviation (dB) | -2.19 ± 1.44 | -10.05<sup>†</sup> | -6.15<sup>*</sup> | -13.35<sup>†</sup> | -6.05<sup>*</sup> |

**Fig. 2.**
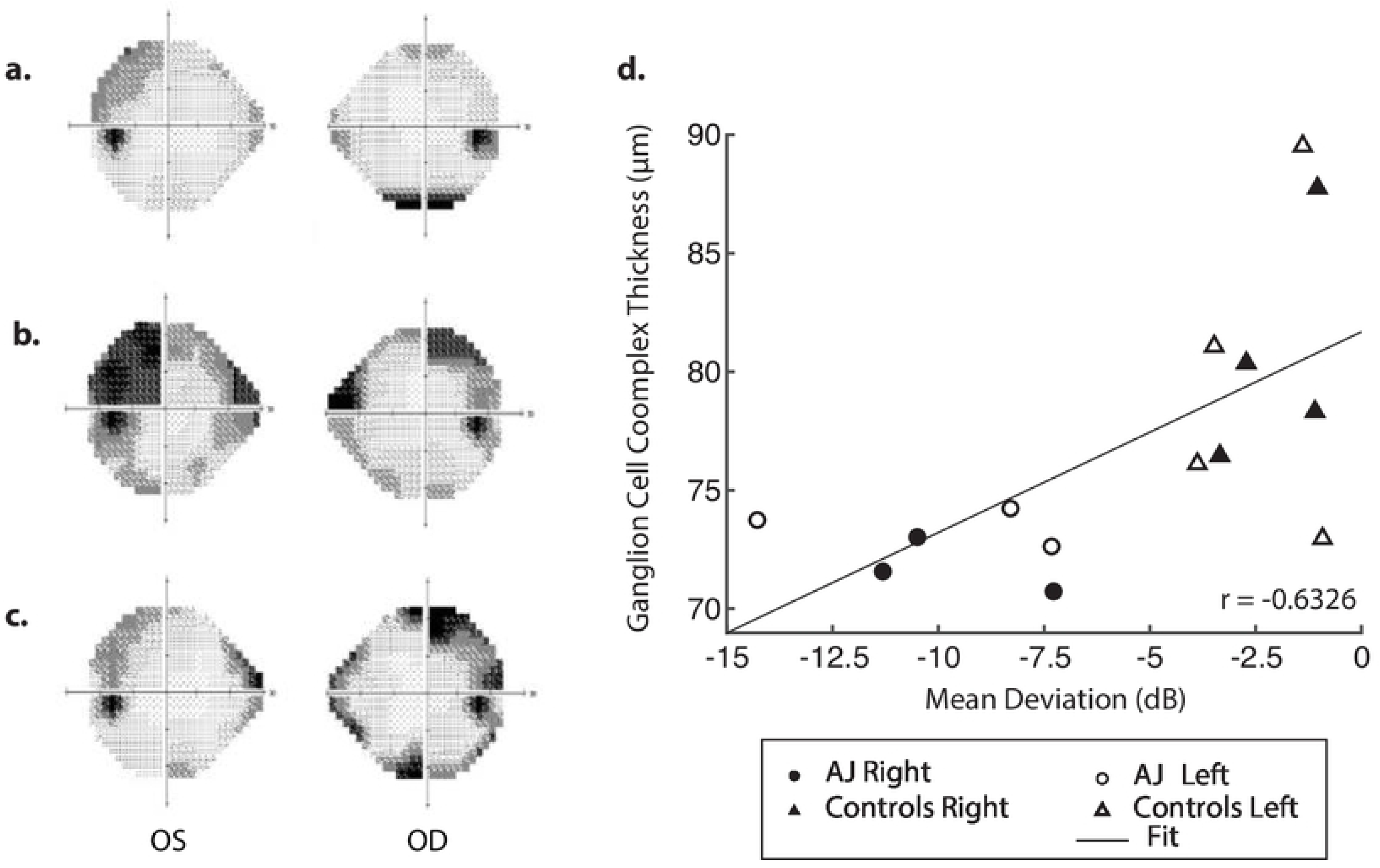
Humphrey 24-2 visual field automated perimetry of AJ at the time of study enrollment (AJ1; panel **a**) illustrating a single temporal defect in OD, mean deviation −6.15 dB, and supratemporal defect in OS, mean deviation −6.57 dB. Following cabergoline use (AJ2; panel **b**), ophthalmologic evaluation was notable for OD supratemporal and nasal defects (mean deviation −9.51 dB) and OS temporal-greater-than-nasal defects (mean deviation −13.35 dB), reflecting worsening of visual function. Further evaluation after final cessation of cabergoline (AJ3; panel **c**), demonstrates subtle worsening of the temporal defect extending infratemporally (mean deviation −9.65) and significantly improved supratemporal and nasal defects (mean deviation −6.05), reflective of overall improvement in visual function. Panel **d** demonstrates a positive correlation between GCC thickness and mean deviation.

#### Retinal Thickness

Optical Coherence Tomography was used to segment the retinal layers in all healthy control participants (n = 8 total eyes) and patient AJ at each time point. There was a trend of reduced retinal thickness across all layers in patient AJ compared with healthy controls (Supplemental Table 4). Notably, this pattern was not significant (all p > 0.05), and thus consistent with values for patient AJ at the border of normal physiologic thickness (75 μm for pRNFL). Prior work indicates that values below the threshold of 75 μm are associated with a low probability of recovery of visual function following injury to the optic tracts (42, 55). Reduced thickness was primarily in the nasal hemiretinas bilaterally (Figure 1E).

GCC data were also correlated with mean deviation, demonstrating a significant relation between retinal thickness and visual function across all study participants (r = 0.63 and p = 0.015; Figure 2D). These data allow us to infer a causal relation between GCC and Humphrey perimetry. It is important however, to note that the stability of retinal thickness measures for patient AJ observed over the duration of the study, suggests that the transient decline in visual function at time point 2 (AJ2) cannot be primarily explained by macrostructural changes within the retina. Raw data for all retinal layers is provided in the supplemental text.

### Diffusion MRI is Sensitive to Alternations in Serum Prolactin Levels

Diffusion MRI was obtained on all study participants, and at three separate time points for patient AJ in tandem with ophthalmologic and laboratory evaluations. At the onset of the study (AJ1), and shortly after initial discontinuation of cabergoline, AJ’s diffusion metrics within the optic tracts were within the range established by the healthy controls sample (Figure 3A-B). This was supported by non-significant Welch’s t-tests (all p > 0.4) comparing AJ’s average diffusion metrics for the optic tracts with those of the control sample (n=10) optic tracts. With decreasing levels of prolactin at the patient’s second visit (AJ2), related to the use of cabergoline, there was a significant increase in average RD compared to the control population (two tailed; p < 0.001) with no change in axial diffusivity (p = 0.52). This difference is apparent in the distribution of voxel-based diffusion measures, as displayed in Figure 3A-B. After following medical advice to again discontinue cabergoline at the third time point (AJ3), diffusion metrics again no longer differed from controls (all p > 0.3). Notably, the pattern relating measures of diffusion in the optic tracts to serum levels of prolactin in patient AJ suggests that higher levels of prolactin are associated with reduced RD. This trend is not present for AD, as demonstrated in Figure 3C-D.

**Fig. 3.**
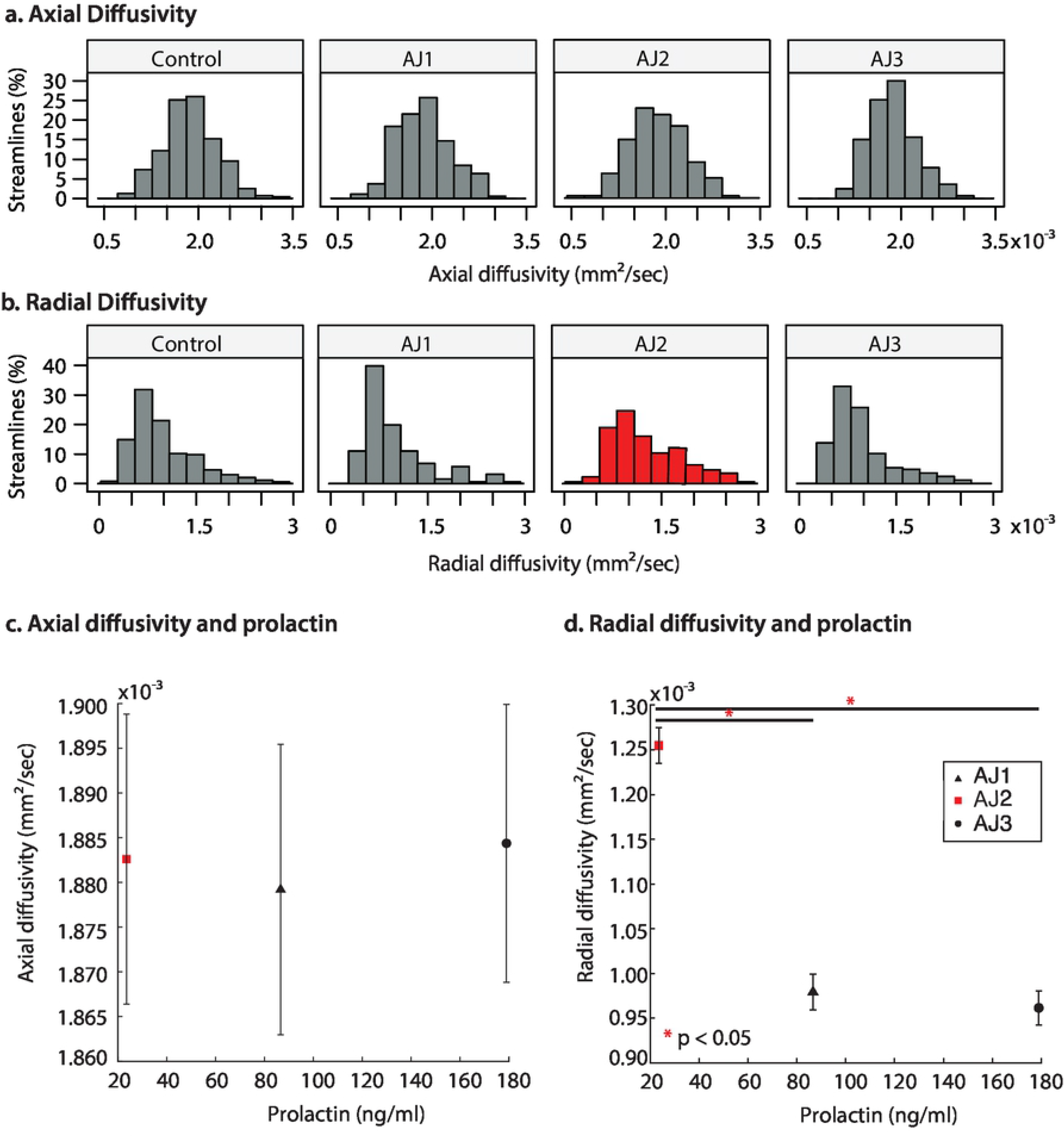
(a-b) Distribution of diffusion MRI metrics for the optic tracts of AJ at three time points and healthy control participants (n=10 total optic tracts). AJ scan 1 (AJ1) was completed after initial discontinuation of cabergoline (high PRL), AJ scan 2 (AJ2) was done after cabergoline was resumed of the patient’s own volition (normal PRL), and AJ scan 3 was done 5 months after final termination of cabergoline treatment (high PRL). Diffusion metrics include axial diffusivity (AD) and radial diffusivity (RD). (c-d) Averaged diffusion metrics for patient AJ as a function of prolactin, demonstrating a negative trend between prolactin and both RD and MD, and a positive trend between prolactin and FA. *p<0.05.

### Diffusion MRI Indices Correlate with Ganglion Cell Complex Thickness

GCC thickness was inversely related to the diffusion MRI index of RD in the optic pathways across all study participants (Pearson correlation, r = −0.6 and p = 0.025), whereas there were no relations for AD and GCC thickness (r = −0.024 and p = 0.94; Figure 4). Some clustering of data points is expected, due to the fact that AJ’s GCC values are expected to be lower than those of the healthy controls. Recognizing that inclusion of patient AJ across time points could potentially introduce unwanted variability in measures of diffusion around a relatively stable GCC thickness, the analyses were repeated for only healthy control participants. Results from those analyses demonstrate the relation remains between measures of diffusion and GCC thickness, for RD (r = −0.69 and p = 0.066), but not between AD and GCC among healthy control participants (r = −0.28 and p = 0.50). These patterns suggest that the identified relations between GCC and diffusion indices represent a baseline structure-function relation between the retina and white matter tract integrity.

**Fig. 4.**
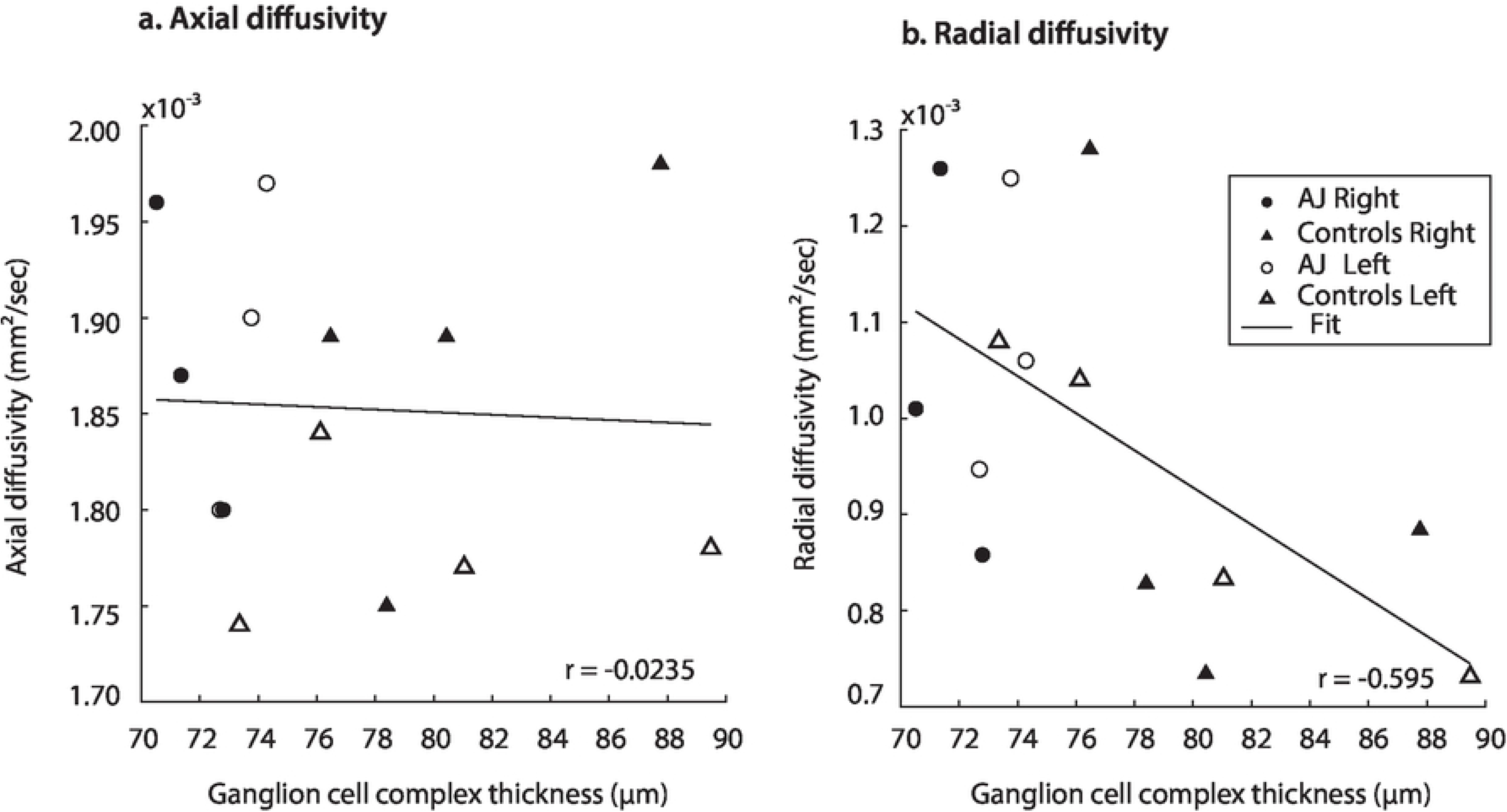
Hemifield GCC thickness measurements are related to indexed diffusion MRI metrics within the corresponding optic tracts. Specifically, significant correlations are identified between GCC and RD (r = −0.595, p = 0.0247). No correlation was identified between GCC and AD (r = −0.0235, p = 0.937). All measures evaluated using linear regression analysis.

### Optic Tract Size does not Correlate with Measures of Diffusion or Hormonal Function

A core finding described above is that diffusion indices of myelination track prolactin levels in patient AJ. In order to rule out the possibility that the observed effects on diffusion indices are derivative of macrostructural changes, such as thinning of the optic tracts secondary to increasing traction on the chiasm, we measured optic tract size for all healthy controls (n = 10 optic tracts) and patient AJ across all time points. The range of right optic tract CSA values was 6.22-9.96 mm^2^ (mean=8.47 mm^2^) for healthy controls and 9.56-10.02 mm^2^ (mean=9.81mm^2^) for patient AJ. In the left optic tract, values ranged from 6.27-11.00 mm^2^ (mean=9.18 mm^2^) for controls and 9.83-10.35mm^2^ (mean=10.08mm^2^) for patient AJ. These data are consistent with previously published studies(54) and there were no right-to-left discrepancies. No statistical differences were identified between optic tract CSA in patient AJ (at any time point) and healthy control participants (p = 0.14, using Welch’s t-test). The tight range of CSA values at each time point demonstrates stability of optic tract size and suggests that increased traction or physical deformation of the optic tract is not the primary factor in the patient’s worsened clinical exam at AJ2. This is further supported by the fact that no correlations were identified between optic tract size and measures of AD, RD, prolactin or GCC, all p > 0.34.

## DISCUSSION

Here, we demonstrate a stereotyped pattern of diffusion MRI changes in response to varying levels of serum prolactin in a rare, isolated, model of nerve traction injury. Specifically, increased RD indexed a significant decrease in serum prolactin. Important to this finding, is that no relations were present between axial diffusivity, levels of prolactin, retinal thickness or visual function. These data confirm, in-vivo, previous observations suggesting a neuroprotective role of prolactin in the human brain (12, 13, 17, 18, 56) and are consistent with the suite of diffusion MRI signals that have been shown to index the integrity of myelin (3, 35–38). Adding to the specificity of these stereotyped diffusion MRI changes is the demonstration of concordant changes in visual ability, indicating disruption of a tightly coupled structure-function relationship between vision and white matter integrity with decreasing serum levels of prolactin. Notably, these patterns were present in the absence of macrostructural changes to the optic tracts, as assessed by CSA and are supported by stable measures of retinal thickness and axial diffusivity. Given the specificity of these findings it is reasonable to infer that serum levels of prolactin causally affect the coupled structure-function relation between white matter integrity (i.e. myelination) and visual ability. Additionally, these data support the use of diffusion MRI as an independent index of the effects of prolactin on white matter in the human brain, both for future observational studies and for future interventional studies.

### Radial diffusivity as an index for glial-neuronal interactions that support myelin health

Prior work has documented a pattern of reduced RD after surgical decompression of the optic chiasm that is disproportionate to changes in AD (5). This pattern is thought to characterize rapid remyelination in the human brain (i.e., days to weeks) following chiasmatic decompression. The current study demonstrates this same pattern of diffusion changes following periods of hyperprolactinemia rather than decompression – and does so in the setting of chronic injury from persistent traction. These data suggest that in the setting of an isolated stretch injury, radial diffusivity indexes the glial-neuronal interactions that support myelin sheath integrity and preservation of function. Key to this conclusion is that despite persistent traction on the optic chiasm throughout the study, white matter injury, as assessed with diffusion MRI, was only made apparent after AJ resumed cabergoline on her own volition – an event that resulted in decreased levels of serum prolactin. This is an important distinction given what is currently known about both neuronal stretch injury (8, 57) and prolactin’s role in neural recovery (13, 17, 18). Once a nerve is stretched, a stereotyped pattern of microstructural changes leads to the development of axonal swelling and correlated changes in the microtubular component of the axonal cytoskeleton, particularly microtubule loss at nodes of Ranvier and at internodal regions (8). More severe injury models describe diffuse patterns of injury with permeabilization of the axolemma and subsequent formation of peri-axonal spaces, myelin inclusions, and reactive axonal swellings culminating in secondary axotomy (57).

There is a growing body of evidence in both in-vitro and in-vivo models demonstrating prolactin’s role in mediating glial-neuronal interactions, including improved astrocyte viability and decreased astrogliosis (12). Critical for inferring microstructural properties from diffusion MRI data is the b-value used during image acquisition: in our case b=1000. At this b-value, diffusion patterns are most sensitive to changes in the extra-axonal compartment (58). Recruitment of oligodendrocyte precursor cells to the injured optic tract can effectively decrease radial diffusivity and is consistent with known properties of prolactin (18). Recovery of both the number and density of microtubules, which has been shown to occur as quickly as 4 hours after an optic nerve stretch injury (8) can also reduce radial diffusivity and represents a potential area of future interest with respect to possible associations with prolactin. These physical changes, which can be measured in the extra-axonal space, likely accompany upregulation of neurotransmission that improves signal conduction and information transfer to the striate cortex. Increased signal conduction alone would be unlikely to provide microstructural changes large enough to be measured at the resolution of diffusion tensor imaging. Data from this study, therefore, add to the argument that prolactin influences a complex network of interactions important for maintaining the health of oligodendrocytes, cytoskeletal structures, and levels of myelination – which would otherwise be susceptible to secondary axonal injury.

### Stability of Retinal Thickness Measures

Recovery after retinal thinning, as measured by OCT varies with pathology (41–43, 45, 55, 59, 60). Optic neuritis patients experience recovery of visual function with increasing VEP amplitudes despite continued loss of pRNFL thickness and significant optic nerve atrophy over 12 months (61). Klistorner and colleagues suggest that this discrepancy is driven by a combination of both remyelination and neural reorganization. When used to measure recovery following nerve decompression (e.g. as treatment for a pituitary macroadenoma), pRNFL and photopic negative response are sensitive to detect changes at three months (41–43, 55). These changes lag behind both improvements in visual function and diffusion MRI indices and are relatively stable at the individual level. In chiasmal compression by pituitary tumors, GCC thickness analysis had greater correlation to mean deviation, a measure of visual defect, than RNFL analysis (62). Additionally, some patients had GCC thinning but no abnormalities in RNFL or mean deviation of visual defect on perimetry testing (63), suggesting increased sensitivity of GCC to injury. As such, this study primarily focused on GCC.

Here, we demonstrate preservation of retinal ganglion cell thickness despite persistent traction injury to the optic nerves, chiasm and tracts over the course of a year, including a transient period (AJ2) of worsening white matter injury and decreased vision. Only one other case with a similar pattern of preserved pRNFL thickness despite recorded deficits on Humphrey perimetry and injury to the optic chiasm has been reported in the literature – a patient with a large, compressive prolactinoma and hyperprolactinemia that was evaluated prior to starting cabergoline (64). While it is possible that the stability of retinal thickness measurements in patient AJ is secondary to exposure to high levels of prolactin – this pattern is complicated by a known delay (approximately 6-8 weeks) in measurable response to pathology, which warrants further investigation. Our findings indicate that radial diffusivity is a sensitive marker that is dynamically modulated in the central nervous system coincident with rapidly changing serum prolactin levels.

### The role of cabergoline

Given that the transient normal prolactin level measured in patient AJ was precipitated by use of cabergoline, it is possible that cabergoline has an independent effect on radial diffusivity and visual function. Chuman and colleagues suggest that cabergoline drug toxicity directly results in vision loss, and that cessation of therapy subsequently leads to vision recovery (22). While plausible, this pattern of delayed vision loss is not typically seen in patients treated with high doses of cabergoline (65), and has not been reported in the absence of empty sella syndrome. Further investigation in animal models relating the impact of cabergoline on D2 receptors in white matter may help to better separate differences between the changes related to prolactin versus those related to dopamine agonists alone. A second hypothesis proposes that in the absence of cabergoline, undetectable tumor regrowth leads to untethering of the of the optic chiasm and a subsequent return of visual function. Two issues arise in this context. First, no patient has been reported to demonstrate improvement in the degree of optic chiasm herniation, at least across all case reports of which we are aware and for which cessation of medical therapy was the treatment strategy (22–27). Second, the amount of deformation of the optic chiasm poorly correlates with visual function, as demonstrated in both compressive pituitary tumor patients (66), and individuals with an incidental finding of primary empty sella syndrome as an anatomic variant (67). As such, we suggest that it is unlikely that an “undetectable” change in tethering is responsible for the significant reduction in radial diffusivity and improved visual function observed with hyperprolactinemia in this study.

## CONCLUSIONS

In summary, we demonstrate in a single patient with empty sella syndrome secondary to dopamine agonist therapy, that increasing serum levels of prolactin correlate with improved visual function and a reduction in radial diffusivity. These data support, in-vivo, a neuro-protective role for prolactin in the injured human brain, confirming work previously conducted in animal models, and establishes radial diffusivity as an important index for tracking the white matter impact of varying levels of serum prolactin. These findings offer a non-invasive means of measuring the effectiveness of novel therapies targeting prolactin as a mediator of neuroprotection in the human brain.

## LIST OF ABBREVIATIONS

dMRI: Diffusion magnetic resonance imaging
OCT: Optical coherence tomography
RNFL: Retinal nerve fiber layer
pRNFL: Peripapillary retinal nerve fiber layer thickness
GCC: Ganglion Cell Complex
IPL: Inner plexiform layer
CST: Macular central subfield
GCL: Ganglion cell layer
OD: Right eye
OS: Left eye
CSA: Cross sectional area
dB: Decibels
AD: Axial diffusivity
RD: Radial diffusivity

## ACKNOWLEDGEMENTS

The authors appreciate Frank Yeh and Arun Venkataraman for their contributions to the development of the diffusion MRI acquisition and analysis pipeline and for their comments on earlier drafts; and to Tod Romo for assistance with computing resources.

## REFERENCES

1. Crowe MJ, Bresnahan JC, Shuman SL, Masters JN, Beattie MS. Apoptosis and delayed degeneration after spinal cord injury in rats and monkeys. Nat Med. 1997;3(1):73–6.

2. Li GL, Farooque M, Holtz A, Olsson Y. Apoptosis of oligodendrocytes occurs for long distances away from the primary injury after compression trauma to rat spinal cord. Acta neuropathologica. 1999;98(5):473–80.

3. Kozlowski P, Raj D, Liu J, Lam C, Yung AC, Tetzlaff W. Characterizing white matter damage in rat spinal cord with quantitative MRI and histology. J Neurotrauma. 2008;25(6):653–76.

4. Totoiu MO, Keirstead HS. Spinal cord injury is accompanied by chronic progressive demyelination. Journal of Comparative Neurology. 2005;486(4):373–83.

5. Paul DA, Gaffin-Cahn E, Hintz EB, Adeclat GJ, Zhu T, Williams ZR, et al. White matter changes linked to visual recovery after nerve decompression. Sci Transl Med. 2014;6(266):266ra173.

6. Rutland JW, Padormo F, Yim CK, Yao A, Arrighi-Allisan A, Huang K-H, et al. Quantitative assessment of secondary white matter injury in the visual pathway by pituitary adenomas: a multimodal study at 7-Tesla MRI. J Neurosurg. 2019;132(2):333–42.

7. Gensert JM, Goldman JE. Endogenous progenitors remyelinate demyelinated axons in the adult CNS. Neuron. 1997;19(1):197–203.

8. Maxwell WL, Graham DI. Loss of axonal microtubules and neurofilaments after stretch-injury to guinea pig optic nerve fibers. J Neurotrauma. 1997;14(9):603–14.

9. DeBoy CA, Zhang J, Dike S, Shats I, Jones M, Reich DS, et al. High resolution diffusion tensor imaging of axonal damage in focal inflammatory and demyelinating lesions in rat spinal cord. Brain. 2007;130(Pt 8):2199–210.

10. Yoles E, Schwartz M. Degeneration of spared axons following partial white matter lesion: implications for optic nerve neuropathies. Exp Neurol. 1998;153(1):1–7.

11. Maxwell WL, Povlishock JT, Graham DL. A mechanistic analysis of nondisruptive axonal injury: a review. J Neurotrauma. 1997;14(7):419–40.

12. Anagnostou I, Reyes-Mendoza J, Morales T. Glial cells as mediators of protective actions of prolactin (PRL) in the CNS. Gen Comp Endocrinol. 2018;265:106–10.

13. Morales T, Lorenson M, Walker AM, Ramos E. Both prolactin (PRL) and a molecular mimic of phosphorylated PRL, S179D-PRL, protect the hippocampus of female rats against excitotoxicity. Neuroscience. 2014;258:211–7.

14. Arnold E, Thebault S, Baeza-Cruz G, Arredondo Zamarripa D, Adan N, Quintanar-Stephano A, et al. The hormone prolactin is a novel, endogenous trophic factor able to regulate reactive glia and to limit retinal degeneration. J Neurosci. 2014;34(5):1868–78.

15. Arnold E, Thebault S, Arona RM, Martinez de la Escalera G, Clapp C. Prolactin mitigates deficiencies of retinal function associated with aging. Neurobiol Aging. 2020;85:38–48.

16. Rivera JC, Aranda J, Riesgo J, Nava G, Thebault S, Lopez-Barrera F, et al. Expression and cellular localization of prolactin and the prolactin receptor in mammalian retina. Exp Eye Res. 2008;86(2):314–21.

17. Patil MJ, Henry MA, Akopian AN. Prolactin receptor in regulation of neuronal excitability and channels. Channels (Austin). 2014;8(3):193–202.

18. Gregg C, Shikar V, Larsen P, Mak G, Chojnacki A, Yong VW, et al. White matter plasticity and enhanced remyelination in the maternal CNS. J Neurosci. 2007;27(8):1812–23.

19. Ezzat S, Asa SL, Couldwell WT, Barr CE, Dodge WE, Vance ML, et al. The prevalence of pituitary adenomas: a systematic review. Cancer. 2004;101(3):613–9.

20. Molitch ME. Diagnosis and Treatment of Pituitary Adenomas: A Review. JAMA. 2017;317(5):516–24.

21. Verhelst J, Abs R, Maiter D, van den Bruel A, Vandeweghe M, Velkeniers B, et al. Cabergoline in the Treatment of Hyperprolactinemia: A Study in 455 Patients. The Journal of Clinical Endocrinology & Metabolism. 1999;84(7):2518–22.

22. Chuman H, Cornblath WT, Trobe JD, Gebarski SS. Delayed visual loss following pergolide treatment of a prolactinoma. J Neuroophthalmol. 2002;22(2):102–6.

23. Taxel P, Waitzman DM, Harrington JF, Jr., Fagan RH, Rothfield NF, Chen HH, et al. Chiasmal herniation as a complication of bromocriptine therapy. J Neuroophthalmol. 1996;16(4):252–7.

24. Dhanwal DK, Sharma AK. Brain and optic chiasmal herniations into sella after cabergoline therapy of giant prolactinoma. Pituitary. 2011;14(4):384–7.

25. Bangash MH, Clarke DB, Holness RO. Brain & chiasmal herniations into sella after medical treatment of prolactinoma. Can J Neurol Sci. 2006;33(2):240–2.

26. Raverot G, Jacob M, Jouanneau E, Delemer B, Vighetto A, Pugeat M, et al. Secondary deterioration of visual field during cabergoline treatment for macroprolactinoma. Clinical Endocrinology. 2009;70(4):588–92.

27. Jones SE, James RA, Hall K, Kendall-Taylor P. Optic chiasmal herniation — an under recognized complication of dopamine agonist therapy for macroprolactinoma. Clinical Endocrinology. 2000;53(4):529–34.

28. Kubo S, Hasegawa H, Inui T, Tominaga S, Yoshimine T. Endonasal endoscopic transsphenoidal chiasmapexy with silicone plates for empty sella syndrome: technical note. Neurol Med Chir (Tokyo). 2005;45:428–32.

29. Gkekas N, Primikiris P, Georgakoulias N. Untethering of herniated left optic nerve after dopamine agonist treatment for giant prolactinoma. Acta Neurochir (Wien). 2013;155(3):495–6.

30. Ishihara E, Toda M, Sasao R, Ozawa H, Saito S, Ogawa K, et al. Endonasal Chiasmapexy Using Autologous Cartilage/Bone for Empty Sella Syndrome After Cabergoline Therapy for Prolactinoma. World Neurosurgery. 2019;121:145–8.

31. Berastegui GRA, Raza SM, Anand VK, Schwartz TH. Endonasal endoscopic transsphenoidal chiasmapexy using a clival cranial base cranioplasty for visual loss from massive empty sella following macroprolactinoma treatment with bromocriptine: case report. J Neurosurg. 2016;124(4):1025–31.

32. Papanastasiou L, Fountoulakis S, Pappa T, Liberopoulos K, Malliopoulos D, Markou A, et al. Brain and optic chiasmal herniation following cabergoline treatment for a giant prolactinoma: wait or intervene? Hormones. 2014;13(2):290–5.

33. Cobb MI-PH, Crowson M, Mintz-Cole R, Husain AM, Berger M, Jang D, et al. Transnasal Transsphenoidal Elevation of Optic Chiasm in Secondary Empty Sella Syndrome Following Prolactinoma Treatment. World Neurosurgery. 2018;112:250–3.

34. Araneda SA, Salgado CM. Visual field defects secondary to cabergoline use in a patient with pituitary tumor. Vision Pan-America. 2015;14(1):2.

35. Xu J, Sun SW, Naismith RT, Snyder AZ, Cross AH, Song SK. Assessing optic nerve pathology with diffusion MRI: from mouse to human. NMR Biomed. 2008;21(9):928–40.

36. Song S-K, Yoshino J, Le TQ, Lin S-J, Sun S-W, Cross AH, et al. Demyelination increases radial diffusivity in corpus callosum of mouse brain. NeuroImage. 2005;26(1): 132–40.

37. Song SK, Sun SW, Ju WK, Lin SJ, Cross AH, Neufeld AH. Diffusion tensor imaging detects and differentiates axon and myelin degeneration in mouse optic nerve after retinal ischemia. NeuroImage. 2003;20(3):1714–22.

38. Song S-K, Sun S-W, Ramsbottom MJ, Chang C, Russell J, Cross AH. Dysmyelination Revealed through MRI as Increased Radial (but Unchanged Axial) Diffusion of Water. NeuroImage. 2002;17(3):1429–36.

39. Smith SA, Williams ZR, Ratchford JN, Newsome SD, Farrell SK, Farrell JA, et al. Diffusion tensor imaging of the optic nerve in multiple sclerosis: association with retinal damage and visual disability. AJNR Am J Neuroradiol. 2011;32(9):1662–8.

40. Engelhorn T, Michelson G, Waerntges S, Otto M, El-Rafei A, Struffert T, et al. Changes of radial diffusivity and fractional anisotropy in the optic nerve and optic radiation of glaucoma patients. ScientificWorldJournal. 2012;2012:849632.

41. Danesh-Meyer HV, Carroll SC, Foroozan R, Savino PJ, Fan J, Jiang Y, et al. Relationship between Retinal Nerve Fiber Layer and Visual Field Sensitivity as Measured by Optical Coherence Tomography in Chiasmal Compression. Investigative ophthalmology & visual science. 2006;47(11):4827–35.

42. Danesh-Meyer HV, Papchenko T, Savino PJ, Law A, Evans J, Gamble GD. In vivo retinal nerve fiber layer thickness measured by optical coherence tomography predicts visual recovery after surgery for parachiasmal tumors. Investigative ophthalmology & visual science. 2008;49(5):1879–85.

43. Moon CH, Hwang SC, Ohn Y-H, Park TK. The Time Course of Visual Field Recovery and Changes of Retinal Ganglion Cells after Optic Chiasmal Decompression. Investigative ophthalmology & visual science. 2011;52(11):7966–73.

44. Crawford JR, Howell DC. Comparing an individual’s test score against norms derived from small samples. The Clinical Neuropsychologist. 1998;12(4):482–6.

45. Schneider CL, Prentiss EK, Busza A, Matmati K, Matmati N, Williams ZR, et al. Survival of retinal ganglion cells after damage to the occipital lobe in humans is activity dependent. Proc Biol Sci. 2019;286(1897):20182733.

46. Kupfer C, Chumbley L, Downer JdC. Quantitative histology of optic nerve, optic tract and lateral geniculate nucleus of man. Journal of anatomy. 1967;101(Pt 3):393.

47. Andersson JL, Skare S, Ashburner J. How to correct susceptibility distortions in spin-echo echo-planar images: application to diffusion tensor imaging. Neuroimage. 2003;20(2):870–88.

48. Andersson JLR, Sotiropoulos SN. An integrated approach to correction for off-resonance effects and subject movement in diffusion MR imaging. Neuroimage. 2016;125:1063–78.

49. Smith SM. Fast robust automated brain extraction. Human Brain Mapping. 2002;17(3): 143–55.

50. Behrens TEJ, Berg HJ, Jbabdi S, Rushworth MFS, Woolrich MW. Probabilistic diffusion tractography with multiple fibre orientations: What can we gain? NeuroImage. 2007;34(1):144–55.

51. Hernandez M, Guerrero GD, Cecilia JM, Garcia JM, Inuggi A, Jbabdi S, et al. Accelerating fibre orientation estimation from diffusion weighted magnetic resonance imaging using GPUs. PLoS One. 2013;8(4):e61892.

52. Hernandez-Fernandez M, Reguly I, Jbabdi S, Giles M, Smith S, Sotiropoulos SN. Using GPUs to accelerate computational diffusion MRI: From microstructure estimation to tractography and connectomes. Neuroimage. 2019;188:598–615.

53. Chernoff BL, Teghipco A, Garcea FE, Sims MH, Paul DA, Tivarus ME, et al. A Role for the Frontal Aslant Tract in Speech Planning: A Neurosurgical Case Study. J Cogn Neurosci. 2018;30(5):752–69.

54. Andrews TJ, Halpern SD, Purves D. Correlated size variations in human visual cortex, lateral geniculate nucleus, and optic tract. The Journal of neuroscience: the official journal of the Society for Neuroscience. 1997;17(8):2859–68.

55. Moon CH, Hwang SC, Kim BT, Ohn YH, Park TK. Visual prognostic value of optical coherence tomography and photopic negative response in chiasmal compression. Investigative ophthalmology & visual science. 2011;52(11):8527–33.

56. Melendez Garcia R, Arredondo Zamarripa D, Arnold E, Ruiz-Herrera X, Noguez Imm R, Baeza Cruz G, et al. Prolactin protects retinal pigment epithelium by inhibiting sirtuin 2-dependent cell death. EBioMedicine. 2016;7:35–49.

57. Povlishock JT, Christman CW. The pathobiology of traumatically induced axonal injury in animals and humans: a review of current thoughts. Journal of neurotrauma. 1995;12(4):555–64.

58. Le Bihan D, Mangin JF, Poupon C, Clark CA, Pappata S, Molko N, et al. Diffusion tensor imaging: concepts and applications. J Magn Reson Imaging. 2001;13(4):534–46.

59. Kerrigan-Baumrind LA, Quigley HA, Pease ME, Kerrigan DF, Mitchell RS. Number of ganglion cells in glaucoma eyes compared with threshold visual field tests in the same persons. Invest Ophthalmol Vis Sci. 2000;41(3):741–8.

60. Trip SA, Schlottmann PG, Jones SJ, Li W-Y, Garway-Heath DF, Thompson AJ, et al. Optic nerve atrophy and retinal nerve fibre layer thinning following optic neuritis: Evidence that axonal loss is a substrate of MRI-detected atrophy. NeuroImage. 2006;31(1):286–93.

61. Klistorner A, Arvind H, Garrick R, Graham SL, Paine M, Yiannikas C. Interrelationship of Optical Coherence Tomography and Multifocal Visual-Evoked Potentials after Optic Neuritis. Investigative Ophthalmology & Visual Science. 2010;51(5):2770–7.

62. Tieger MG, Hedges TR, 3rd, Ho J, Erlich-Malona NK, Vuong LN, Athappilly GK, et al. Ganglion Cell Complex Loss in Chiasmal Compression by Brain Tumors. J Neuroophthalmol. 2017;37(1):7–12.

63. Blanch RJ, Micieli JA, Oyesiku NM, Newman NJ, Biousse V. Optical coherence tomography retinal ganglion cell complex analysis for the detection of early chiasmal compression. Pituitary. 2018;21(5):515–23.

64. Lukewich MK, Micieli JA. Chronic chiasmal compression and persistent visual field defect without detectable changes in optical coherence tomography of the macular ganglion cell complex. Am J Ophthalmol Case Rep. 2019;16:100533.

65. Ono M, Miki N, Kawamata T, Makino R, Amano K, Seki T, et al. Prospective Study of High-Dose Cabergoline Treatment of Prolactinomas in 150 Patients. The Journal of Clinical Endocrinology & Metabolism. 2008;93(12):4721–7.

66. Porciatti V, Ciavarella P, Ghiggi MR, D’Angelo V, Padovano S, Grifa M, et al. Losses of hemifield contrast sensitivity in patients with pituitary adenoma and normal visual acuity and visual field. Clinical Neurophysiology. 1999;110(5):876–86.

67. Griffiths PG, Dayan M, Coulthard A. Primary empty sella: cause of visual failure or chance association? Eye (Lond). 1998;12 (Pt 6):905–6.

